# Pattern Of Some Inflammatory Mediators Among Adult Patients With Bronchial Asthma On Corticosteroids Attending Aminu Kano Teaching Hospital, Kano, Nigeria

**DOI:** 10.1101/2022.08.08.503171

**Authors:** H Saidu, M Ismail, U Lawal, H Mannir, MY Gwarzo, LD Rogo, A Ibrahim, N Garba, SB Danladi, IA Aliyu, JA Bala, IS Yahya

## Abstract

**Background:** Bronchial asthma in adults is typified by lingering allergic inflammation associated with elevation in the levels of certain acute phase reactants and indicators of mast cell activation. This study investigated the effect of corticosteroid treatment on the levels of C reactive protein (CRP), serum baseline tryptase (sBT), erythrocyte sedimentation rate (ESR) and granulocyte monocyte colony stimulating factor (GM-CSF) among asthmatics.

**Method:** Forty five adult patients with bronchial asthma on treatment with inhaled corticosteroids were enrolled. Forty five blood donors were used as control. Serum levels of CRP, sBT, ESR and GM-CSF were measured using sandwich ELISA.

**Result:** The GM-CSF, CRP, ESR and sBT were significantly elevated among asthmatics on treatment compared to normal healthy control. Significant difference in the level of GM-CSF, ESR and CRP was observed between asthmatics with mild and moderate disease severity.

**Conclusion:** Treatment with inhaled corticosteroids doesn’t restore the levels of GM-CSF, CRP, sBT and ESR to normalcy among asthmatics.

## Introduction

The major aims of asthma management are to minimize and prevent asthma symptoms, improve quality of life, cut the rate and severity of asthma exacerbations, and reverse airflow obstruction (Ohta et al., 2011). This is supposedly achieved through a judicious use of a cocktail of drugs and life style changes. This approach supposedly reverses the pathologic cycle of the disease and prevents deterioration in the quality of life (Wang *et al*., 2016). However, the success of this approach is disappointing in Nigeria, as acute attacks and long term complications are not uncommon among asthmatics in the country (Ozoh *et al*., 2019). The reasons for this failure may be multifactorial; poor response to drug treatment due to disease heterogeneity, poor adherence to treatment protocol and lack of heed to experts’ advice by the patients and unavailability of resources to adequately diagnose and monitor the disease are some of the identified factors (Desalu et al., 2011). The heterogeneity in the immunobiological mechanisms of asthma contributes immensely to variation in disease phenotype. This reiterates the need for biomarkers by allergists and respiratory physicians good enough to predict prognosis and monitor treatment success (Tiotiu, 2018). The commonest used agents in the management of asthma are the beta 2 receptor agonists and corticosteroids (Ohta et al., 2011). These drugs reverse the vicious cycles of smooth muscle contraction in the airways and minimize local inflammation. The inflammatory reaction is made evident serologically by elevation of certain acute phase reactants like C-reactive protein (CRP) and erythrocyte sedimentation rate (ESR). The role of GM-CSF in allergic inflammation and autoimmune diseases is increasingly appreciated (Shiomi and Usui, 2015). Moreover, the characteristic eosinophil activation in asthma is enhanced by the release of granulocyte monocyte colony stimulating factor (GM-CSF) by activated TH2 cell, macrophages, airway epithelium and fibroblasts (Shiomi and Usui, 2015). Mast cells are believed to occupy a special position in the pathogenesis of asthma, basically through the release of vasoactive early and late phase mediators such as histamine and cysteinyl leukotriense. They also produce cytokines that skew the immune response towards a Th2 phenotype (Varricchi *et al*., 2019). Activation of mast cells is easily demonstrated by the measurement of serum baseline tryptase(Schroeder *et al*., 2016). It is preferred to histamine due to its stability and longer half-life (Schroeder *et al*., 2016). However, the exact nature of the effect of treatment with corticosteroids on these allergic inflammatory biomarkers is not adequately explored in Nigeria (Ozoh et al., 2019). Thus this study investigated the pattern of serum baseline tryptase (sBT), GM-CSF, CRP and ESR among bronchial asthma patients on corticosteroids thus paving the way for their use in patient monitoring.

## Method

This is a comparative cross sectional hospital based study, conducted in Aminu Kano teaching hospital Kano of Nigeria. Forty five adult patients (18-70 years of age) with bronchial asthma were enrolled into the study. The participants have asthma of either mild or moderate grade and were on maintenance treatment with low dose inhaled corticosteroids for at least one month to the time of enrollment. None of the patients was having acute exacerbation as at the time of recruitment. Patients with other respiratory illnesses were excluded. Patients with HIV infection and other inflammatory diseases like hypertension and diabetes were also excluded. The control were apparently healthy blood donors devoid of the commonly screened transfusion transmissible infections in the region (HIV, HCV, HBV and Syphilis). They have no current history of respiratory diseases. Ethical permission from the management of Aminu Kano teaching hospital Kano was obtained (AKTH/MAC/SUB/12A/P-3/VI/2699) and informed consent of the participants was sought before enrollment.

Five milliliters of blood sample was collected from all eligible participants, 1.6 ml was dispensed immediately into a sample bottle containing 0.4ml of sodium citrate and the remaining 3.4 ml was dispensed into a plain container. Serum was obtained from the clotted sample in the plain container and used for the measurement of GM-CSF, CRP and sBT. The sodium citrated sample was used for the measurement of ESR. Sandwich ELISA method based kits obtained from monobind Lake Forest USA and Melsin Medical Co limited, Chunchang China were used to measure the levels of GM-CSF, CRP and sBT. The ESR was determined using the Westergren method.

## Result

The levels of serum GM-CSF, sBT and CRP in the study group were found to be 101.38 ± 20.46ng/l, 19.69±5.39 ng/l and 3.65±0.58ug/ml respectively. The levels of the parameters among the control group were found to be 77.45 ± 77.00ng/l, 14.88±6.86 ng/l and 2.74 ± 0.77 ug/ml respectively. There was statistically significant difference in the concentrations of the measured parameters between the study and control group, p= 0.047, < 0.001, < 0.001 and < 0.001 respectively. Similar result was obtained in the ESR between the two groups viz 22.03 + 21.14 and 6.88 3.82, p value = < 0.001. Significant difference in the level of GM-CSF, CRP and ESR was observed between participants with mild and those with moderately severe asthma: 91.20 ±25.16 ng/l and 129.45 ±34.32 ng/l, 3.65 ±0.58ug/ml and 3.89±0.77ug/ml and 19.03± 16.15mm/hr and 24.88 ±18.22mm/hr; p values of 0.020, 0.025 and 0.021 respectively.

**Table 1.** Medical history of the participants.

| Characteristics | Frequency | Percentage (%) |
| --- | --- | --- |
| **Severity** |  |  |
| Mild persistent | 20 | 44.4 |
| Moderate persistent | 25 | 55.6 |
| **Drugs** |  |  |
| Albuterol | 45 | 100% |
| Inhaled Corticosteroid |  |  |

**Table 2:** Comparative analysis of inflammatory biomarkers between study and the control group.

|  | Test group (n=45) | Control group (n=45) | T statistic | P value |
| --- | --- | --- | --- | --- |
| **GMCSF\*(ng/l)** | 101.38 ± 20.46 | 77.45 ± 77.00 | 1.675 | 0.047 |
| **sBT#(ng/l)** | 19.69±5.39 | 14.88±6.86 | 3.873 | <0.001 |
| **CRP#(µg/ml)** | 3.65±0.58 | 2.74±0.77 | 3.985 | <0.001 |
| **ESR\*(mm/hr)** | 22.03±21.14 | 6.88±3.82 | 4.322 | <0.001 |

\* summarized as median ±IQR and ManWhitney U test was used for significance test # summarized as mean ±SD and unpaired t test was used for significance test

**Table 3:** Association between disease severity and inflammatory biomarkers.

|  | Mild (n=20) | Moderate (n=25) | T statistic | P value |
| --- | --- | --- | --- | --- |
| **GMCSF\*(pg/ml)** | 91.20 ±25.16 | 129.45 ±34.32 | 2.975 | 0.020 |
| **sBT#(pg/ml)** | 19.39±3.97 | 20.43±8.15 | 1.201 | >0.005 |
| **CRP#(µg/ml)** | 3.65±0.58 | 3.89±0.77 | 2.321 | 0.025 |
| **ESR\*(mm/hr)** | 19.03±16.15 | 24.88±18.22 | 2.872 | 0.021 |

\* summarized as median ±IQR, ManWhitney U test was used for significance test. # summarized as mean ±SD, unpaired t test was used for significance test

## Discussion

The serum level of granulocyte-monocyte colony stimulating factor is only slightly elevated among the asthma patients. It is also more elevated among patients with moderate disease compared to those with mild presentation. The cytokine GM-CSF is incriminated in the pathogenesis of bronchial asthma through its ability to recruit and maintaining the survival of eosinophils and other immunocytes (Park *et al*., 1998). Seldom is attention paid to measurement of serum level of GM-CSF in asthma; this is due to the understanding that asthma is a disease typified by local inflammation in the airways with less possibility of significant variations in the systemic levels of various molecules coordinating the inflammation (Federico et al., 2006, Saha et al., 2009). However in this study, we are able to report some differences in the serum level of the cytokine GM-CSF between asthmatics and normal healthy control. This difference is also manifested across various grades of the disease according to our findings. The latter finding is in agreement with what was reported by Federico et al., 2006, that elevated GM-CSF level is found in the sputa of asthmatics with severe disease. We were not able to include treatment naïve asthmatics in this study to further control the effect of treatment on the biomarkers. Moreover, the patients were limited to mild and moderate categories of the disease. Nonetheless, from this study it is clear that asthmatics on corticosteroids do not have a fully restored GM-CSF levels compared to the normal healthy control. Whether this is due to poor disease control or inherent inability of the drugs to lower the release of the cytokine is yet to be fully elucidated in the region. However, Sullivan *et al*, 1996 reported insignificant elevation in serum GM-CSF among asthma patients. They however reported raised levels in the bronchoalveolar lavage fluid (BAF) derived eosinophils following allergen challenge (Sullivan *et al*, 1996). In this study we were not able to measure GM-CSF in BAF.

The serum baseline tryptase level was found to be elevated among the cases compared to the control. This is in line with what was reported by Goa et al., (2016); when they compared the pattern of sBT between asthmatics and normal healthy individuals. Abdel Gawad et al., (2017) also reported elevated level of the mast cell activation marker among asthmatics compared to healthy control. It is obvious from this study that evidence of mast cell activation persist even among asthmatics on corticosteroids. This study could not find a link between sBT and asthma disease severity, we hypothesized this to be the effect of corticosteroid treatment the patients are receiving; corticosteroids were found to inhibit mast cell recruitment and the late phase reaction (Grossman and Jensen, 1984, Busse, 1984 and Williams, 2018). Previous studies have indicated association between sBT and disease severity among participants with atopic asthma and its potential to be used as a predictor of asthma control (Gao *et al*., 2016).

We observed that the acute phase reactants CRP and ESR are significantly elevated among asthmatics compared to the normal healthy control. This is in agreement to what was reported in many other studies (Monadi *et al*., 2016, Kasayama *et al*., 2009, Deraz *et al*., 2012). The observed elevation in the level of CRP is however not dramatic, as it is still within the normal range of 2-10ug/ml (Gurhal R and Jialal, 2020). However, the erythrocyte sedimentation rate is elevated above normal values expected of the age range of the asthmatics despite treatment with corticosteroids i.e <12mm/h at 23^0^C (Bates, 2017). Though ESR and CRP are all acute phase reactants, it is not surprising however that the latter is not elevated to above normal range. ESR is determined by a lot of biological factors ranging from the relative concentrations of other acute phase reactants like fibrinogen to the proportion of red cells in the peripheral blood (Vanessa *et al*., 2019). Though corticosteroids are known to have pleiotropic effects on various organ systems (Williams, 2018); from this study it is clear that acute phase response is still ongoing among the asthmatics compared to normal controls. Subsequent studies may have to among other things include an additional group of untreated cases and also explore the relationship between drug dosage and serum level of the biomarkers to further appreciate their kinetics.

## Conclusion

Serum CRP, GM-CSF, sBT and ESR are elevated among asthmatics on inhaled corticosteroid.

## Author’s contribution

Saidu H conceived the idea supervised the work and drafted the initial manuscript, Ismail M, Mannir H and Lawal U performed the experiments and statistical analysis. Gwarzo MY, Rogo LD, Ibrahim A, Garba N, Danladi SB, Aliyu IA, Bala JA, Yahya IS proofread and approved the final draft

